# Genome-wide maps of highly-similar intrachromosomal repeats that mediate ectopic recombination in three human genome assemblies

**DOI:** 10.1101/2024.01.29.577884

**Authors:** Luis Fernandez-Luna, Carlos Aguilar-Perez, Christopher M. Grochowski, Michele Mehaffey, Claudia M.B. Carvalho, Claudia Gonzaga-Jauregui

## Abstract

Repeated sequences spread throughout the genome play important roles in shaping the structure of chromosomes and facilitating the generation of new genomic variation. Through a variety of mechanisms, repeats are involved in generating structural rearrangements such as deletions, duplications, inversions, and translocations, which can have the potential to impact human health. Despite their significance, repetitive regions including tandem repeats, transposable elements, segmental duplications, and low-copy repeats remain a challenge to characterize due to technological limitations inherent to many sequencing methodologies.

We performed genome-wide analyses and comparisons of direct and inverted repeated sequences in the latest available human genome reference assemblies including GRCh37 and GRCh38 and the most recent telomere-to-telomere alternate assembly (T2T-CHM13). Overall, the composition and distribution of direct and inverted repeats identified remains similar among the three assemblies but we observed an increase in the number of repeated sequences detected in the T2T-CHM13 assembly versus the reference assemblies. As expected, there is an enrichment of repetitive regions in the short arms of acrocentric chromosomes, which had been previously unresolved in the human genome reference assemblies. We cross-referenced the identified repeats with protein-coding genes across the genome to identify those at risk for being involved in genomic disorders. We observed that certain gene categories, such as olfactory receptors and immune response genes, are enriched among those impacted by repeated sequences likely contributing to human diversity and adaptation.

Through this analysis, we have produced a catalogue of direct and inversely oriented repeated sequences across the currently three most widely used human genome assemblies. Bioinformatic analyses of these repeats and their contribution to genome architecture can reveal regions that are most susceptible to genomic instability. Understanding how the architectural genomic features of repeat pairs such as their homology, size and distance can lead to complex genomic rearrangement formation can provide further insights into the molecular mechanisms leading to genomic disorders and genome evolution.

**Author summary:** This study focused on the characterization of intrachromosomal repeated sequences in the human genome that can play important roles in shaping chromosome structure and generating new genomic variation in three human genome assemblies. We observed an increase in the number of repeated sequence pairs detected in the most recent telomere-to-telomere alternate assembly (T2T-CHM13) compared to the reference assemblies (GRCh37 and GRCh38). We observed an enrichment of repeats in the T2T-CHM13 acrocentric chromosomes, which had been previously unresolved. Importantly, our study provides a catalogue of direct and inverted repeated sequences across three commonly used human genome assemblies, which can aid in the understanding of genomic architecture instability, evolution, and disorders. Our analyses provide insights into repetitive regions in the human genome that may contribute to complex genomic rearrangements

## Introduction

Repeated sequences spread throughout the genome play a role in genomic instability, structural variant complexity, and the generation of complex genomic rearrangements that may contribute to disease burden (1,2). The formation of new structural variants (SVs), including copy-number variants (CNVs), inversions, and translocations through genomic rearrangements is largely influenced by the local genomic architecture and partially defined by the presence and local density of repetitive sequences (3–11). Defining the molecular features of these repeated sequences including their size, homology, orientation, and genome-wide distribution can help identify substrates to predict rearrangement-prone regions. Such an approach led to the discovery of new genomic disorders (12,13), insights into somatic and germline organismal mutational processes (5,14), and better understanding of mechanisms contributing to genomic rearrangements and genome evolution (15,16).

Studying the genomic features of repeated sequences can help identify unstable genomic regions containing inverted and direct repeat pairs that can act as substrates for recombination or rearrangement events. Non-allelic homologous recombination (NAHR) (17) and replication-based mechanisms (RBMs) such as break-induced replication (BIR) (18), Fork Stalling and Template Switching (FoSTeS) (19) and Microhomology Mediated Break-Induced Replication (MMBIR) (20) utilize highly identical genomic tracts of varying sizes to mediate double-stranded break (DSB) repair (2,21). NAHR can cause large interchromosomal and intrachromosomal deletions, duplications, translocations, and inversions depending on variables such as size, orientation, the degree of identity, and the distance between low-copy repeats (LCRs) or other repetitive elements that serve as substrates (21,22). The crossovers between directly oriented or inverted LCRs that flank distinctive genomic regions driven by NAHR events generally result in recurrent rearrangements that are responsible for most of the well-known and characterized genomic disorders (2,17,23–25). In contrast, non-recurrent genomic rearrangements which can include complex rearrangements produced by microhomology-mediated mechanisms (MMBIR/FoSTeS) have been identified in other less common disorders (26–28). RBMs promote strand invasion and subsequent DNA synthesis resulting in non-recurrent genomic rearrangements which underlie the molecular cause of >70 genetic syndromes (26–29). Data from well-studied genomic disorders such as *MECP2* duplication syndrome (MDS, MIM #300260) and Pelizaeus-Merzbacher disease (PMD, MIM #312080) indicate that directly oriented paralogous low-copy repeats (LCRs) are associated with the formation of simple tandem duplications (19,30–32) and deletions, likely because they facilitate formation of higher-order structures prone to DSBs (33). In contrast, inverse paralogous low-copy repeat pairs seem to have a distinct role of mediating repair of DSBs through BIR which may lead to a high incidence of complex genomic rearrangement formation in loci flanked by those repeats (28,34–36).

The improvement of long-read genomic sequencing technologies in recent years is now facilitating the sequencing and assembly of new whole genomes with great accuracy independently of using a reference assembly. Most recently, a gapless alternate human genome assembly was produced by the Telomere-to-Telomere Consortium adding and assembling 8% of the human genome that had remained elusive in the reference assembly (37). The newly characterized regions in the T2T-CHM13v2 (T2T-CHM13) assembly provide a more comprehensive view of the organization and structure of human repetitive regions that are mainly composed of tandemly arrayed repeats, segmental duplications, and complex repeats in pericentromeric and subtelomeric regions (38). Among previously unresolved regions, the T2T-CHM13 assembly also provides sequence and context for the ribosomal RNA gene clusters in the short arms of acrocentric chromosomes 13, 14, 15, 21, and 22, which had remained unassembled in the GRCh37 and GRCh38 human genome reference assemblies (39).

Previous genome-wide analyses of repetitive elements focused on identifying segmental duplications have used different computational and experimental approaches. However, given the specific characteristics of segmental duplications, these analyses do not present a complete landscape of repeated elements in the human genome (40,41). Other studies focused on the genome-wide identification of paralogous low-copy repeats in the GRCh37 (hg19) assembly used parameters tailored to prioritize identification of NAHR substrates (34,42). As the study of genomic rearrangements has evolved and the understanding of the mechanisms involved and their corresponding substrates has increased, we have improved and expanded the parameters to be considered in the identification of repeated sequences that can contribute to genomic rearrangements. Here we report genome-wide bioinformatic analyses and comparisons of repeated sequences in the three latest and currently most widely used human genome assemblies, GRCh37, GRCh38 and T2T-CHM13. We utilized more comprehensive parameters based on years of experimental data to analyze and identify direct and inverted repeated sequences that can serve as potential substrates for genomic rearrangements. We compare the characteristics and genome-wide distribution of these repeats among the analyzed assemblies. These analyses provide a more comprehensive profile of the organization, distribution, and structure of repeated sequences across the human genome and between assemblies. We focused on the intersection of the identified repeats with protein-coding genes, known disease genes, and previously reported human structural variants. Finally, we discuss their potential role in mediating genome instability and evolution.

## Results

### Genome-wide landscape of identical repeats in the human genome

Through our bioinformatic analyses, we obtained datasets for direct and inverted intrachromosomal repeated sequence pairs for each of the current human genome assemblies, the two latest human genome reference assemblies, GRCh37 (hg19) and GRCh38 (hg38), and the most recent telomere-to-telomere alternate assembly, T2T-CHM13. A total of 570829, 573085 and 585604 repeated sequence pairs in direct orientation were found in the GRCh37, GRCh38, and T2T-CHM13 assemblies, respectively. Similarly, we identified 611838, 612089 and 627791 inverted repeat pairs for each of the assemblies analyzed (Table 1). Given our parameters, the percent identity between pairs of repeats ranged from 80% to 100% for direct and inverted repeats across assemblies. The median percent identity for all identified repeat pairs in both direct and inverted orientations across the three assemblies was approximately 84.2%. The distance between pairs of repeated sequences had a median of 30-31 Mb across assemblies (Figure 1, Table 1). The size of the identified repeats ranged from the minimum size parameter of 200 bp to several hundred base pairs in both direct and inverted orientation across the three assemblies, with the largest repeat expanding 1.6 Mb identified in the T2T-CHM13 assembly (Table 1, Figure 1). Notably, over half of the repeats, approximately 62-64% across the three different assemblies, had a length below 1 kilobase (Table 1, Figure 2, Supplementary Table 1, Supplementary Figure 1). The major peak observed in the three assemblies between 200bp and 400bp corresponds primarily to *Alu* elements, which have an average length of 300bp (Figure 2) and other SINE elements. The peak at 1kb corresponds with the greatest number of segmental duplications with a continuous distribution beyond 1kb. Another major peak can be observed in the 6kb to 7kb size range representing primarily LINE elements in the human genome (Figure 2). Statistics on the main features of the identified repeats in each dataset are reported in Table 1.

**Figure 1.**
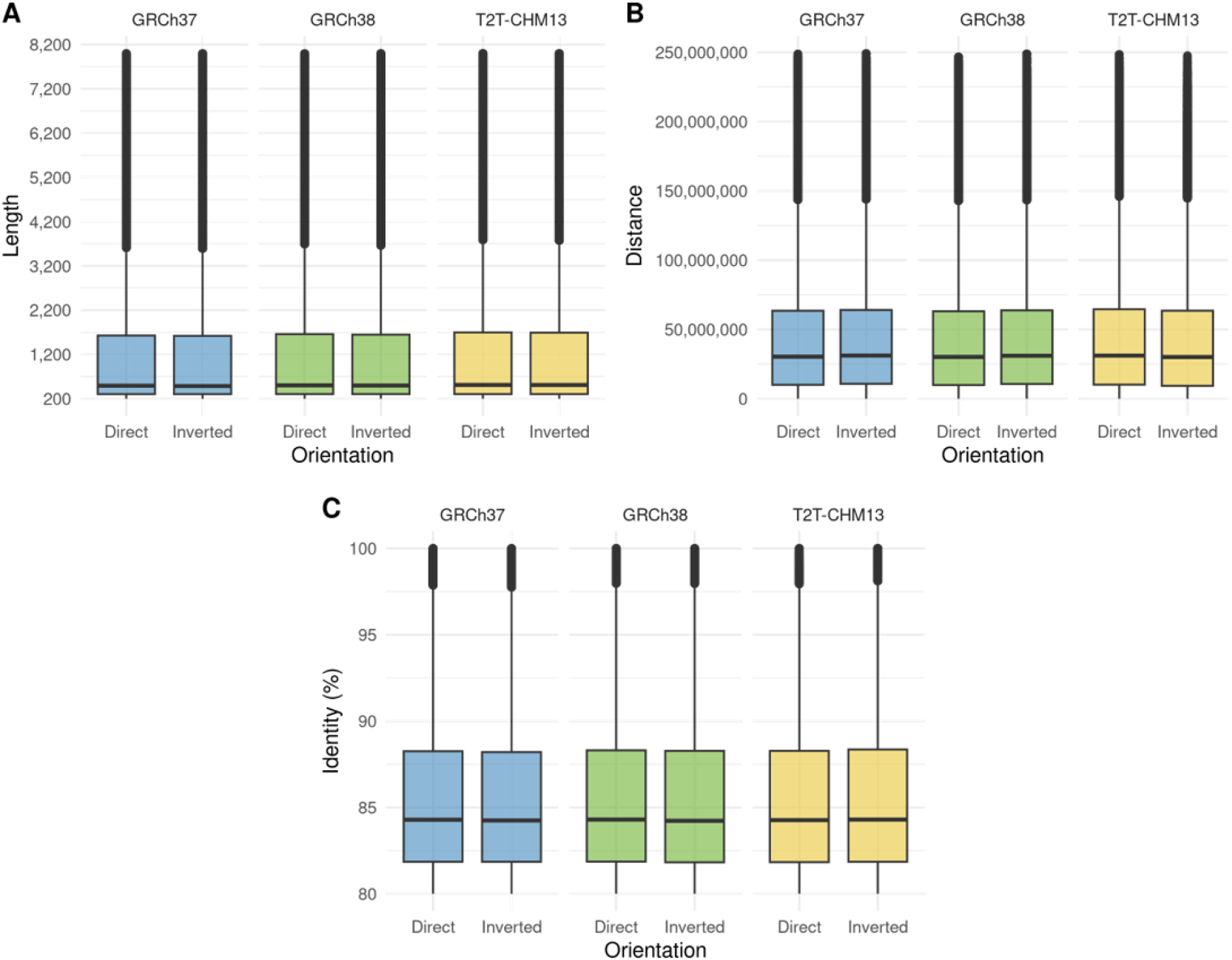
General statistics of the identified direct and inverted repeated sequences across the three human genome assemblies (GRCh37, GRCh38, and T2T-CHM13) analyzed. The descriptive statistics for the identified consolidated identical repeats are similar across the three assemblies. A) Length distribution. B) Distance distribution between pairs of repeats. C) Pairwise percent identity distribution.

**Figure 2.**
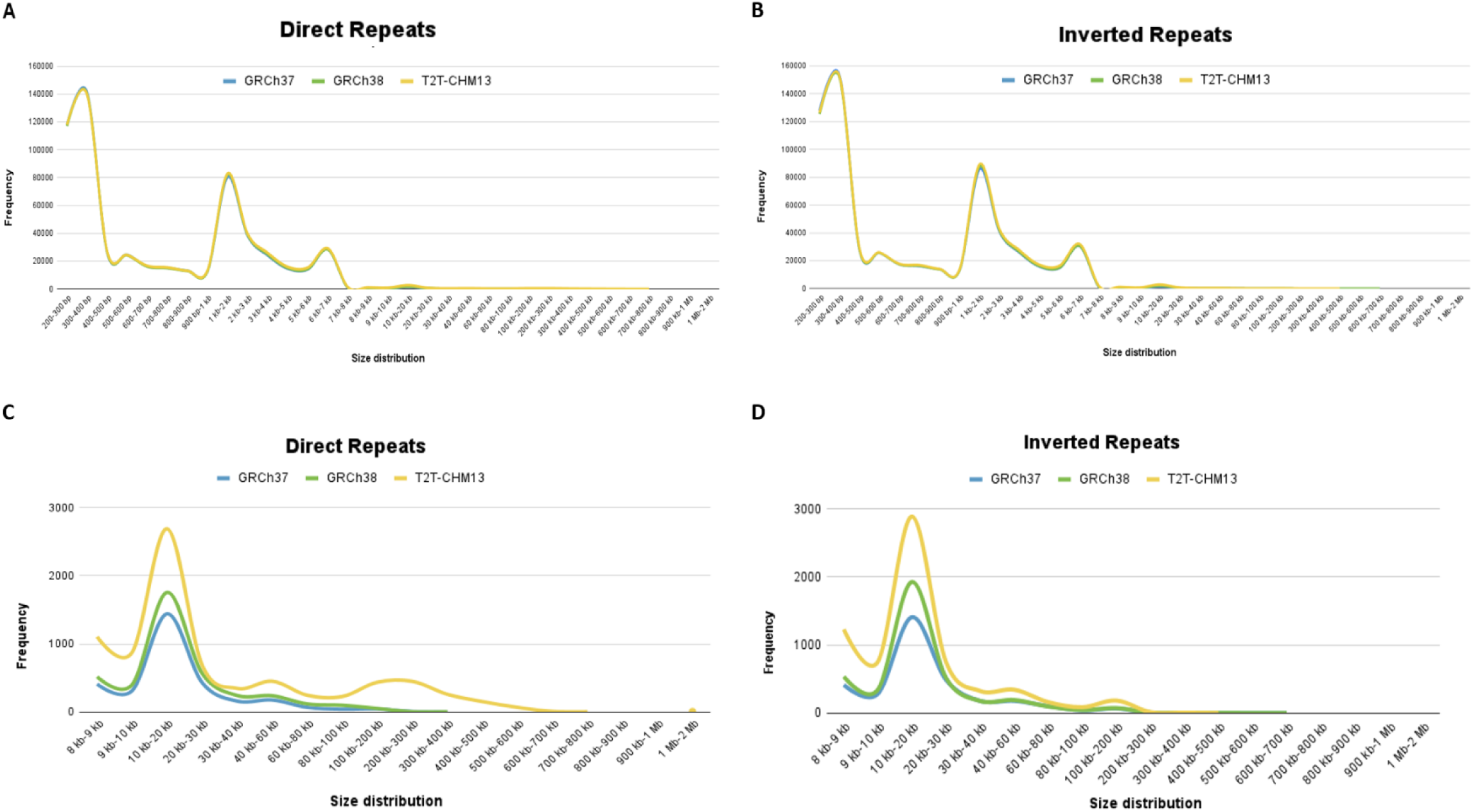
Size distribution of direct and inverted repeated sequences across the genome in the analyzed assemblies (GRCh37, GRCh38, and T2T-CHM13). The overall size distribution of direct and inverted repeats across the genome was observed to be very similar for the three assemblies. A) Size distribution of direct repeats across assemblies. B) Size distribution of inverted repeats across assemblies. C) Examining size distribution of direct repeats beyond the 8 kb size range. Note a slight increase of direct repeats in T2T-CHM13 compared to the two reference assemblies D) Examining size distribution of inverted repeats beyond the 8 kb size. Note a slight increase of inverted repeats in T2T-CHM13 compared to the two reference assemblies.

**Table 1.** Overall statistics of identified direct and inverted repeated sequences.

|  |  | Direct |  |  | Inverted |  |  |
| --- | --- | --- | --- | --- | --- | --- | --- |
|  |  | GRCh37 | GRCh38 | T2T-CHM13 | GRCh37 | GRCh38 | T2T-CHM13 |
| **Total repeat pairs (N)** |  | 570,829 | 573,085 | 585,604 | 611,838 | 612,089 | 627,791 |
| **Total repeat pairs (bp,%)** |  | 320525555,<br>(10.35%) | 328261336,<br>(10.63%) | 391552384,<br>(12.56%) | 328497932,<br>(10.61%) | 336565558,<br>(10.90%) | 365728379,<br>(11.73%) |
| **Length (bp)** | *Min-Max* | 200-395595 | 200-395596 | 200-1663487 | 200-647494 | 200-647495 | 200-495481 |
|  | *Median* | 500 | 509 | 525 | 469 | 505 | 520 |
|  | *Mean* | 1450 | 1503 | 2263 | 1455 | 1483 | 1612 |
|  | *<1kb (N, %)* | 367079<br>(64.31%) | 364350<br>(63.58%) | 367493<br>(62.75%) | 394368<br>(64.46%) | 390579<br>(63.81%) | 395079<br>(62.93%) |
| **Identity (%)** | *Min-Max* | 80-100 | 80-100 | 80-100 | 80-100 | 80-100 | 80-100 |
|  | *Median* | 84.29 | 84.3 | 84.3 | 84.25 | 84.23 | 84.27 |
|  | *Mean* | 85.48 | 85.5 | 85.52 | 85.45 | 85.46 | 85.47 |
| **Distance (bp)** | *Min-Max* | -14-248756443 | -8-246623862 | -8-247651277 | -9995-249229690 | -2-248935491 | 163-248381454 |
|  | *Median* | 30263091 | 30098648 | 30016196 | 31119448 | 30978896 | 31094072 |
|  | *Mean* | 42739049 | 42431293 | 42353751 | 43275746 | 43075283 | 43149712 |

We observed a similar genome-wide distribution of direct and inverted repeated sequence pairs across chromosomes for the three assemblies, except for the short arms of the acrocentric chromosomes (13, 14, 15, 21, and 22) in the T2T-CHM13 assembly (Figure 3, Figure 4). As expected, we were able to detect repeated sequences in these regions due to the availability of new genomic sequence data in the T2T-CHM13 assembly for these chromosomal arms compared to the reference assemblies. In addition to the short arms of acrocentric chromosomes, we detected an increased number of repeated sequences in chromosome Y of the T2T-CHM13 assembly, probably due to a better resolution provided by long-read sequencing of the repeated nature of this chromosome.

**Figure 3.**
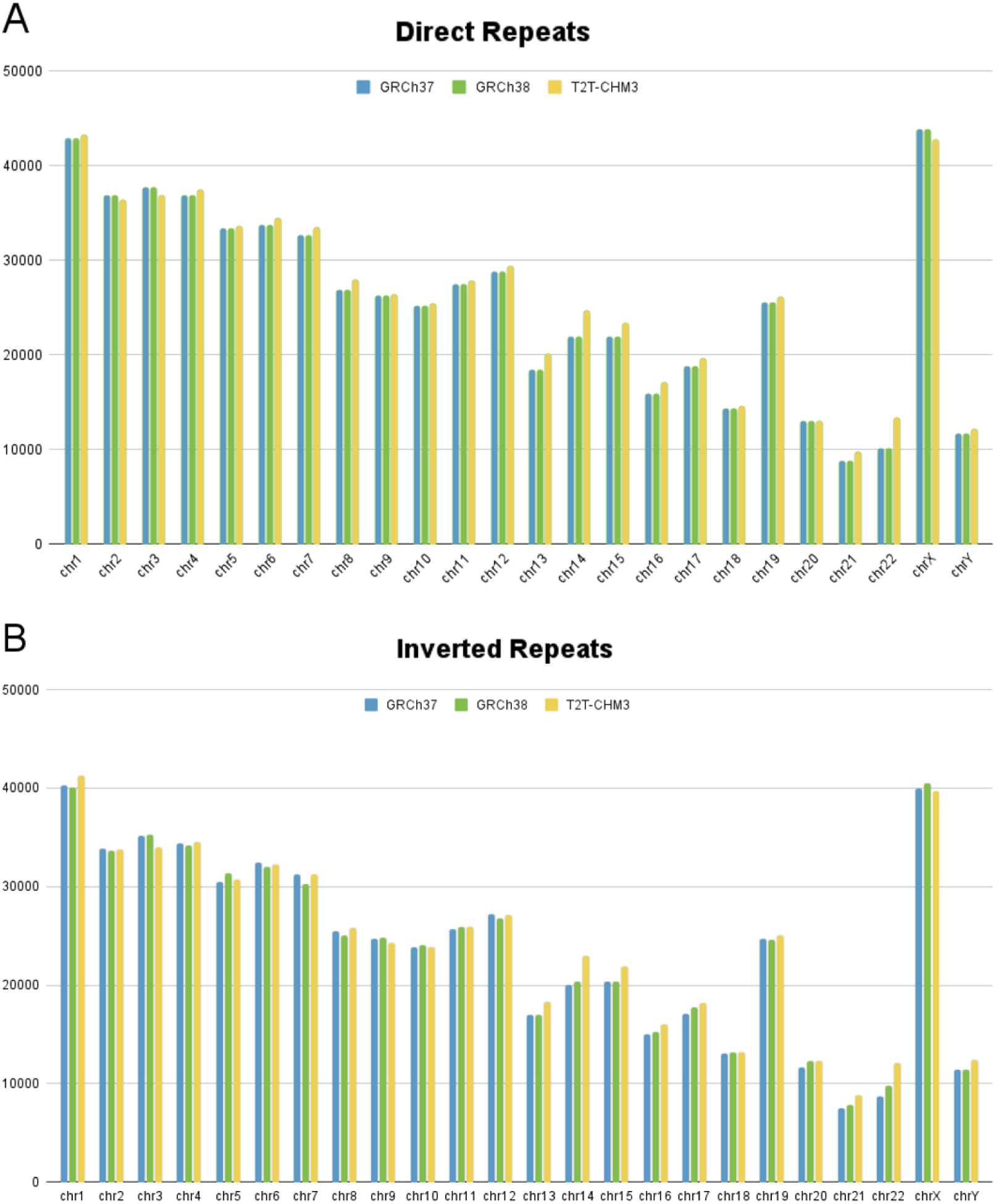
Per chromosome distribution of direct and inverted repeated sequences across the genome in the analyzed assemblies (GRCh37, GRCh38, and T2T-CHM13). The overall distribution of direct and inverted repeats across the genome was observed to be very similar for the three assemblies we studied except for a higher number of repeats detected in the acrocentric chromosomes (13, 14, 15, 21 and 22) and the Y chromosome of the T2T-CHM13 assembly. A) Per chromosome distribution of direct repeats across assemblies. B) Per chromosome distribution of inverted repeats across assemblies.

**Figure 4.**
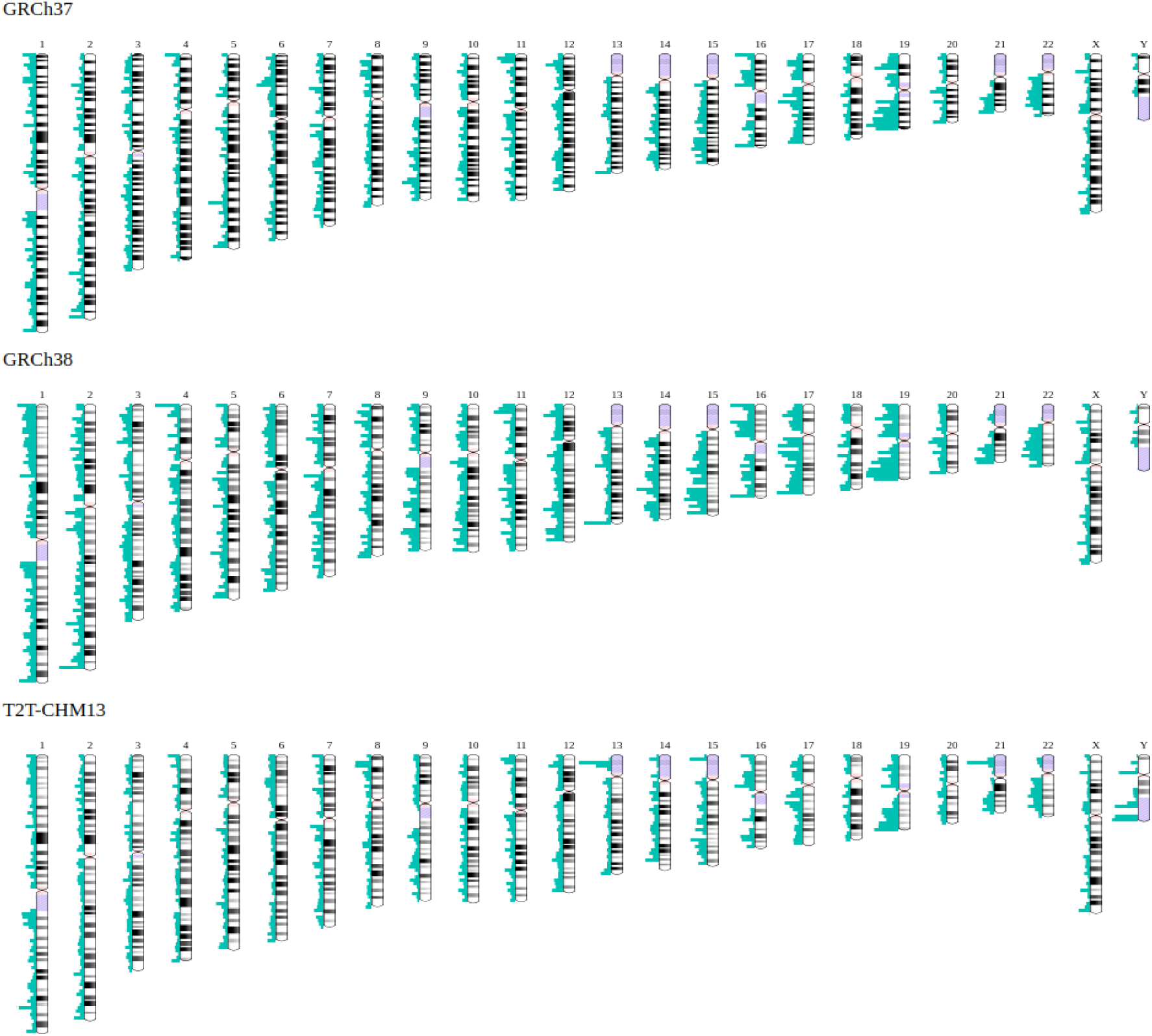
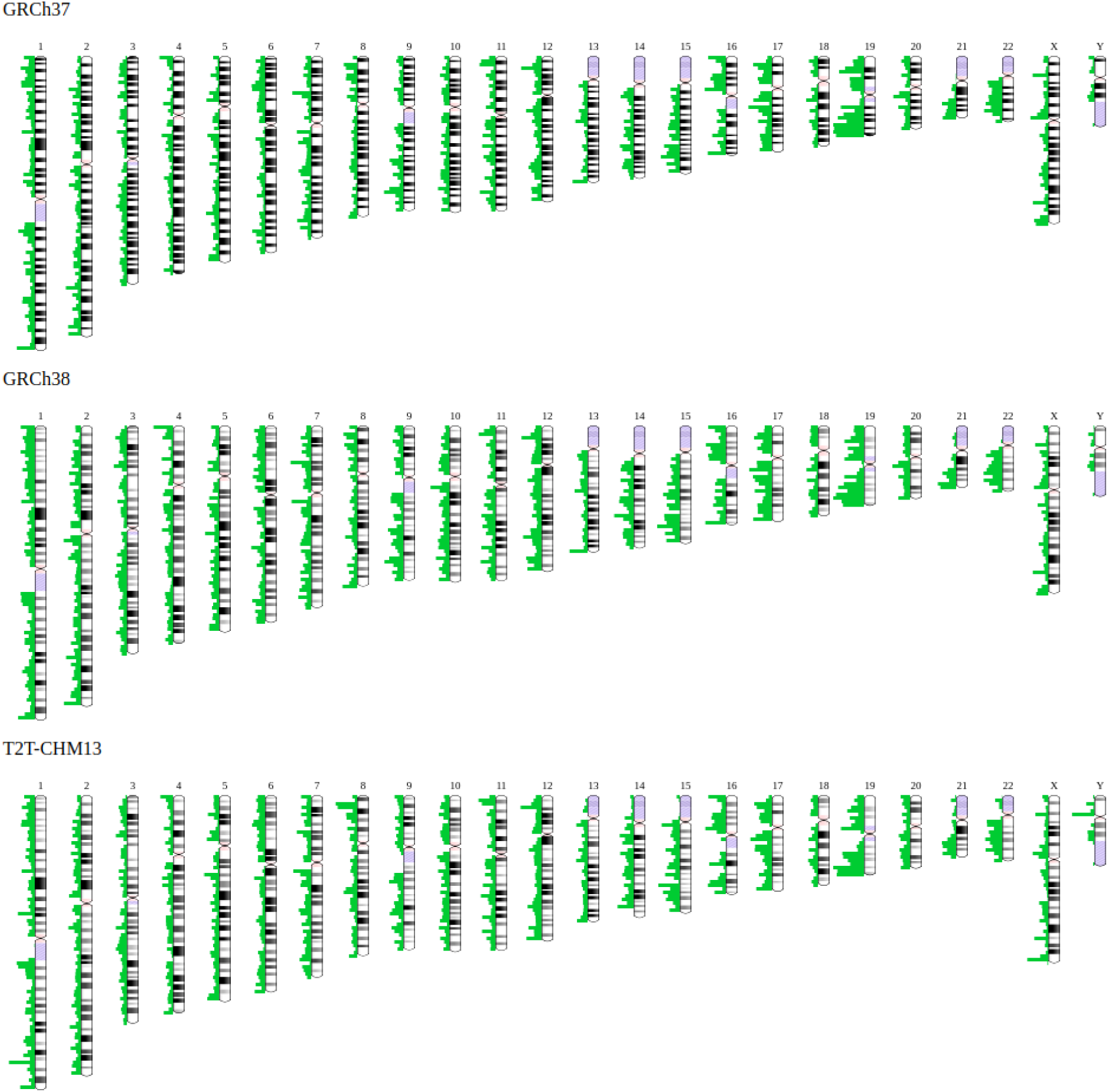
Ideograms of the human chromosomes showing the distribution of direct and inverted repeated sequences identified in this study across three human genome assemblies analyzed (GRCh37, GRCh38, and T2T-CHM13). A) Genome-wide distribution of direct repeats across human chromosomes in the top panel, in blue. B) Genome-wide distribution of inverted repeats across human chromosomes in the lower panel, in green. Note the overrepresentation of repeated sequences in the short arms of acrocentric chromosomes (13, 14, 15, 21 and 22) and the Y chromosome in the T2T-CHM13 assembly versus the two reference assembly versions for both direct and inverted repeats.

The identified direct and inverted repeated sequences are distributed throughout the genome (Figure 3, Figure 4). The percent of base pairs encompassed by direct and inverted repeats considered separately in the reference assemblies ranges from 10.3% to 10.9% (GRCh37 direct 10.35%; inverted 10.61% and GRCh38 direct 10.63%; inverted 10.90%). This percentage of base pairs encompassed by repeated sequences is greater for T2T-CHM13 (direct 12.56% and inverted 11.73%) (Table 1). The overall non-overlapping base pairs of both directed and inverted repeat sets for each assembly are 13.12% (GRCh37), 13.50% (GRCh38), and 15.6% (T2T-CHM13) of the corresponding genome sequences (Supplementary Table 1). This increase in percent of base pairs covered likely reflects the progress in improving and polishing the sequence of the different assemblies using novel sequencing technologies.

### Composition and annotation of direct and inverted repeats across the genome

Genomic rearrangements in the genome can contribute to human disease by directly affecting the structure or normal dosage of genes, or indirectly by altering the appropriate regulation of relevant genes through positional effects or deletion/duplication of regulatory elements (25). The majority of recurrent genomic rearrangements derive from recombination events involving relatively large (>1kb) and highly similar (>90%) repetitive elements such as low-copy repeats (LCRs) or segmental duplications (SDs) through NAHR (2,12,25). Other smaller and more divergent repetitive elements, such as short interspersed nuclear elements (SINEs) and long interspersed nuclear elements (LINEs), may also act as recombination substrates for genomic rearrangements, primarily through microhomology-mediated mechanisms (MMBIR/FoSTeS). More specifically, *Alu* elements, a common subclass of SINEs representing 11% of the genome, have been observed to be involved in *Alu*/*Alu*-mediated rearrangements (43,44).

In order to better characterize the identified repeated sequences datasets and gain further insights into their impact on known genomic elements, we cross-referenced these with cataloged repetitive elements and known protein-coding genes in the human genome.

In the T2T-CHM13 assembly, larger proportions of direct and inverted repeat sequence pairs (10.5% and 12.4% respectively) were observed overlapping segmental duplications compared to the reference assemblies (6.2% and 6.7% for direct and inverted repeats in GRCh37; 7.6% and 8.2% for direct and inverted repeats in GRCh38, respectively) due to the increased number of SDs mapped and annotated in T2T-CHM13 and previously overlooked in the heterochromatic regions of the reference assemblies. Other types of repeats such as LINEs, SINEs and *Alu*s were similarly distributed across the genome in all three assemblies, except, as expected, for the newly assembled regions in T2T-CHM13 not present in the two reference assemblies (Table 2, Figure 5).

**Figure 5.**
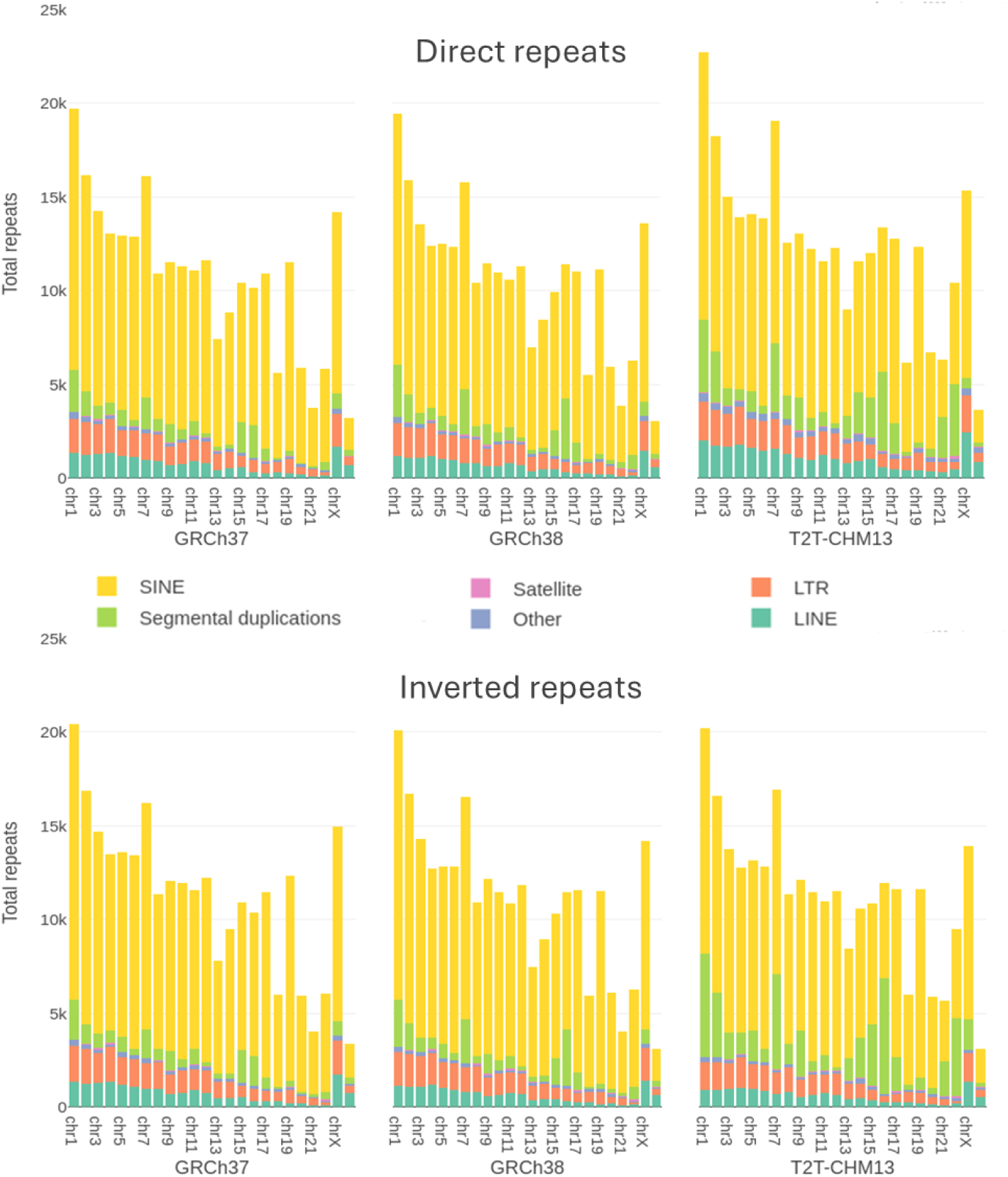
Overlap of identified repeated sequences with known repetitive genomic elements across human genome assemblies. A large fraction of repeat pairs was observed overlapping with SDs, LINEs, and satellite repeats in T2T-CHM13 compared to the reference assemblies. Overlap with other types of repetitive genomic elements was similar across the different assemblies. All autosomes, 1-22, X and Y chromosomes are represented. A) Repeated genomic elements that overlap with the directly repeated sequences from our analysis. B) Repeated genomic elements that overlap with inverted repeated sequences from our analysis.

**Table 2.** Repeat pairs overlapping repetitive elements across the three genome assemblies. The presence of repetitive elements is similar across all assemblies. The percentages are given based on the matching fraction of our consolidated direct and inverted repeats respectively with repetitive genomic elements.

|  | GRCh37 |  | GRCh38 |  | T2T-CHM13 |  |
| --- | --- | --- | --- | --- | --- | --- |
| Repeat type | Direct | Inverted | Direct | Inverted | Direct | Inverted |
| SINE | 194063<br>(74.85%) | 205439<br>(75.90%) | 189389<br>(74.66%) | 200233<br>(75.79%) | 198184<br>(66.43%) | 199981<br>(73.27%) |
| LINE | 18316<br>(7.06%) | 18193<br>(6.72%) | 15688<br>(6.18%) | 15566<br>(5.89%) | 26694<br>(8.94%) | 15505<br>(5.68%) |
| LTR | 25000<br>(9.64%) | 25579<br>(9.45%) | 23757<br>(9.36%) | 24377<br>(9.22%) | 28431<br>(9.53%) | 23977<br>(8.78%) |
| SDs | 17470<br>(6.73%) | 17046<br>(6.29%) | 20874<br>(8.22%) | 20099<br>(7.66%) | 37099<br>(12.43%) | 28764<br>(10.53%) |
| Satellite | 115 (0.04%) | 100 (0.03%) | 94 (0.03%) | 92 (0.03%) | 480 (0.16%) | 288 (0.10%) |
| Other | 4273 (1.64%) | 4302 (1.58%) | 3852 (1.51%) | 3815 (1.44%) | 7404 (2.43%) | 4418 (1.61%) |

### Gene overlap

We observed 933, 970 and 914 protein-coding genes overlapped by direct repeats; while 834, 872, and 847 overlapped with inverted repeats in GRCh37, GRCh38 and T2T-CHM13, respectively (Table 3). Similarly, 1872, 1803 and 1796 protein-coding genes were flanked by direct repeat pairs, while 1413, 1574 and 1428 genes were flanked by inverted repeated sequences within a range of 100 kb upstream or downstream in the corresponding assemblies (Table 3). Overall, a total of 3663, 3774, and 3652 non-redundant genes can be potentially affected by rearrangements of the repeated sequences we have identified respectively in the three assemblies (Supplementary table 2). Next, given the known impact of copy-number variants and genomic rearrangements on dosage-sensitive genes, we looked for genes deemed to be dosage-sensitive according to the study by Dong et al (45) among the protein-coding genes impacted by the identified repeats. A majority of potentially affected genes are considered dosage-sensitive according to the analyses of Dong et al. Furthermore, we observed a higher number of dosage-sensitive genes flanked by repeated sequence pairs rather than overlapping with them. In the GRCh37, GRCh38, and T2T-CHM13 assemblies, a total of 1783, 1834, and 1778 potentially dosage-sensitive genes were flanked by repeated sequence pairs, respectively. Conversely, in each of the assemblies, there were 561, 573, and 550 potentially dosage-sensitive genes that were overlapped by identified repeated sequences (Figure 6).

**Figure 6.**
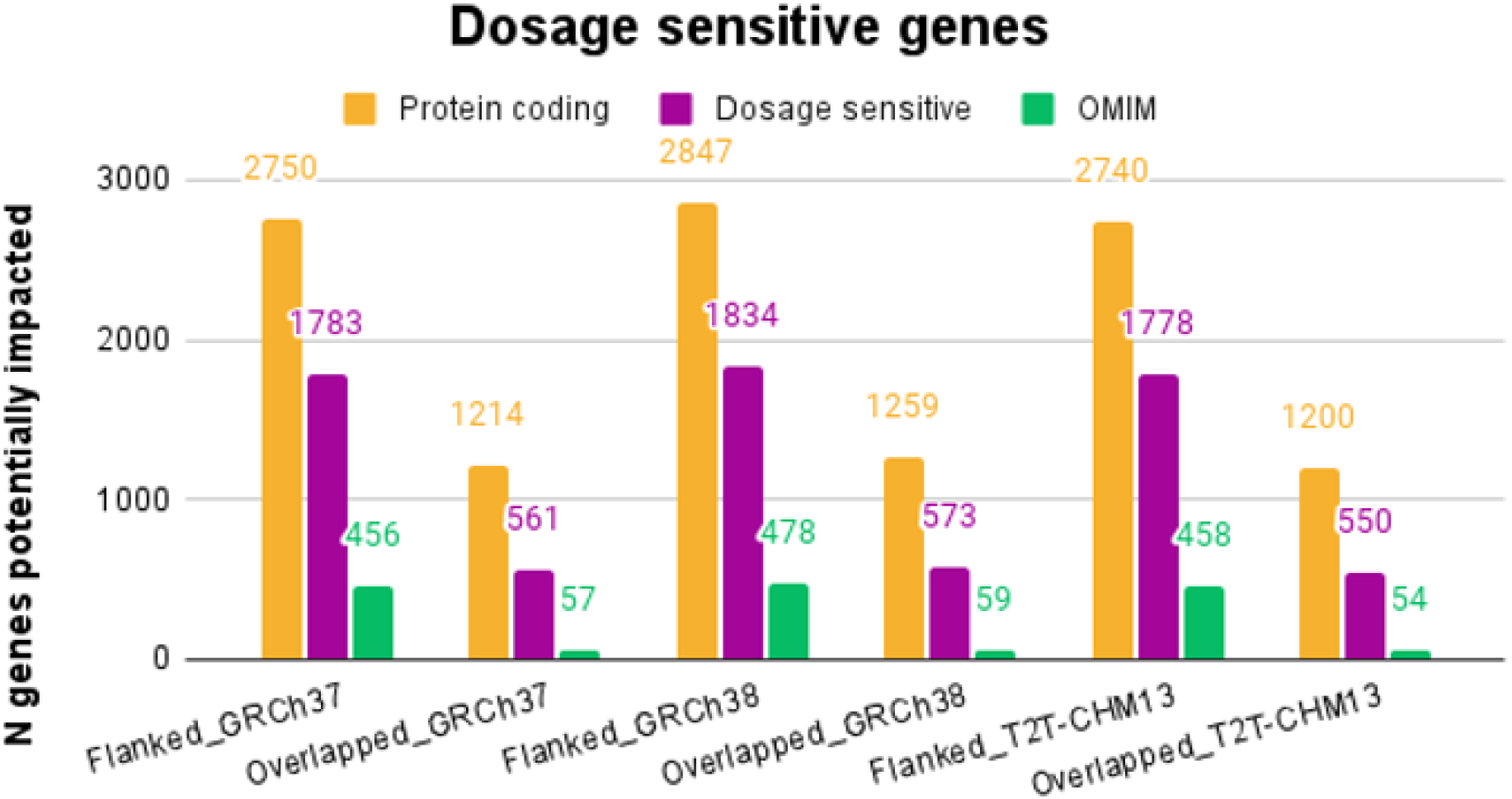
Protein coding genes potentially impacted by the identified repeated sequences. Number of protein-coding genes, dosage-sensitive genes and OMIM genes associated with diseases, found to be overlapped or flanked by identified direct and inverted repeats across the three assemblies.

**Table 3.** Protein-coding genes potentially impacted by the identified consolidated repeats datasets. Number of protein-coding genes, including genes associated with diseases, found to be overlapped or flanked by identified direct and inverted repeats.

|  | GRCh37 |  | GRCh38 |  | T2T-CHM13 |  |
| --- | --- | --- | --- | --- | --- | --- |
| Protein coding genes | Direct | Inverted | Direct | Inverted | Direct | Inverted |
| Overlapped | 933 | 834 | 970 | 872 | 914 | 847 |
| Flanked | 1872 | 1413 | 1803 | 1574 | 1796 | 1428 |
| OMIM_genes |  |  |  |  |  |  |
| Overlapped | 46 | 33 | 49 | 36 | 47 | 35 |
| Flanked | 371 | 259 | 347 | 289 | 348 | 262 |

To analyze the functional implications of the genes potentially affected by genomic rearrangements, a gene ontology (GO) enrichment analysis was performed. This analysis revealed that across all three assemblies, the main enriched classes were related to the olfactory system, such as the detection of chemical stimulus (GO:0050911), sensory perception of smell (GO:0007608), and GPCR signaling (GO:0007186) (Figure 7). This is consistent with the well-known genetic and functional variability observed in the olfactory receptor (OR) gene family, and the contribution of CNVs and genomic arrangements in their inter-individual and cross-population variation (46–48). In addition to the olfactory system, we also identified enrichment for genes associated with immune response and metabolic processes among the top 10 GO terms across the three assemblies (Figure 7).

**Figure 7.**
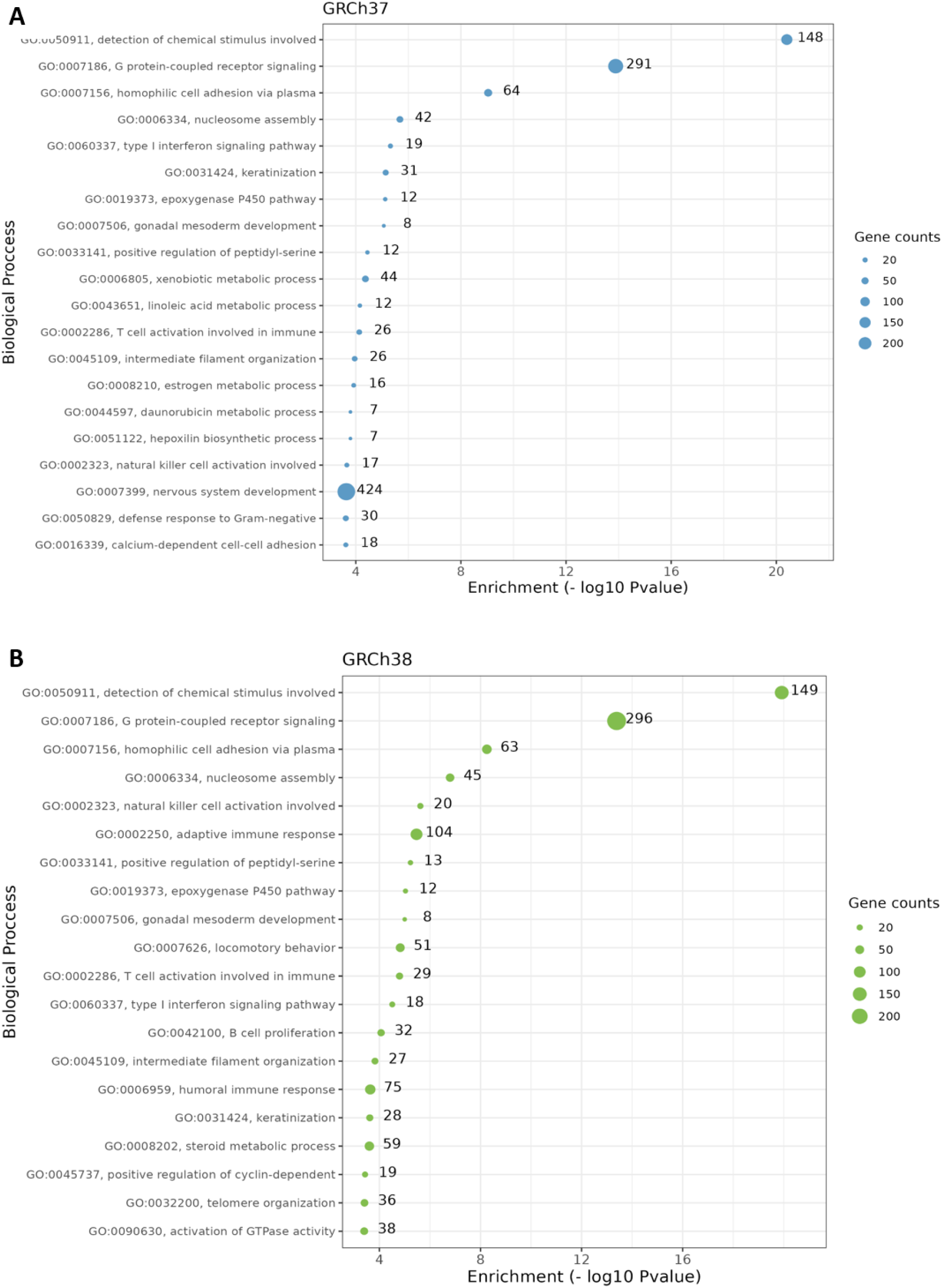
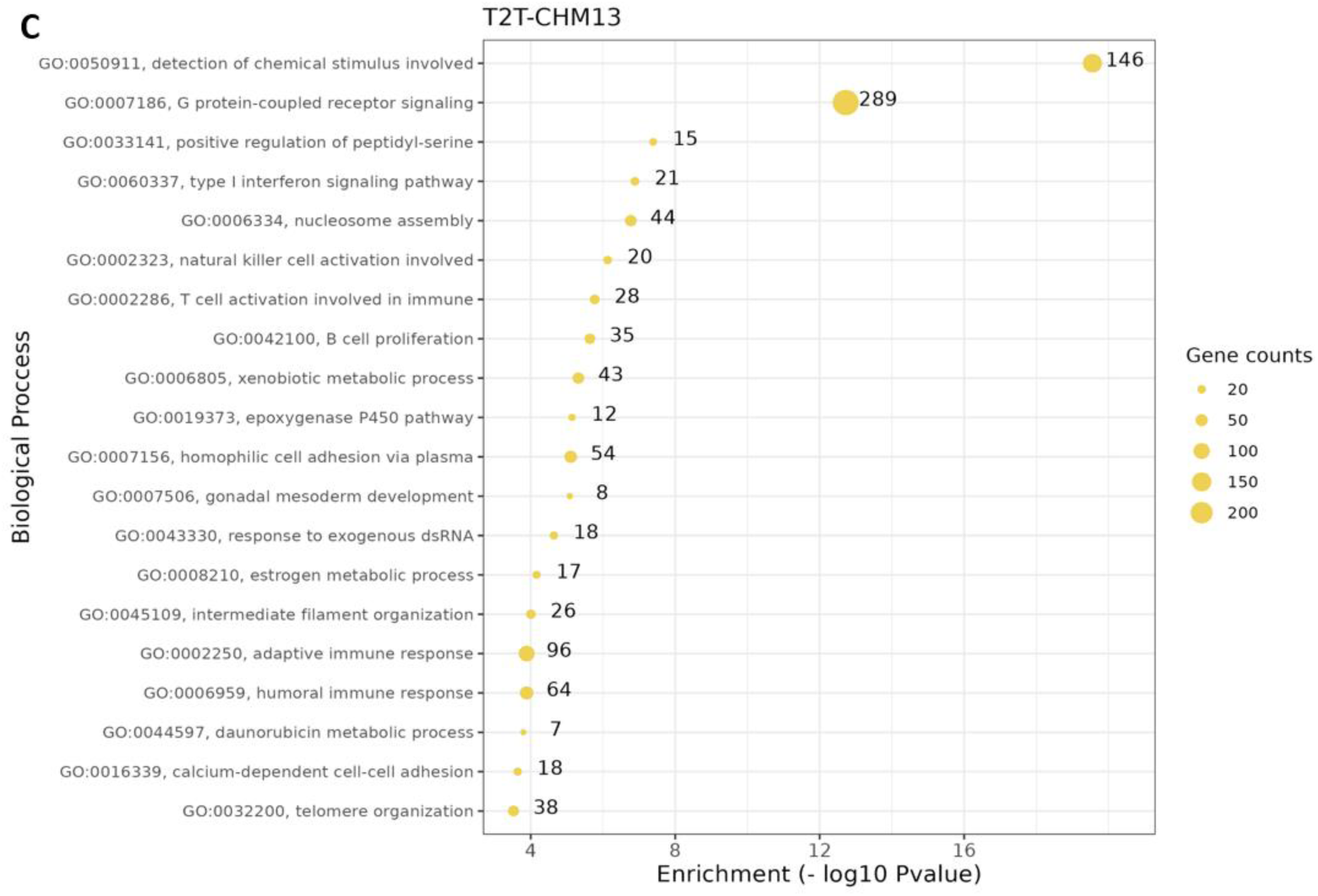
Gene ontology enrichment analysis of genes potentially impacted by repeated sequences. Our analyses showed that genes related to the olfactory system, G protein-coupled receptor signaling, and a few immune and metabolic processes were enriched in regions overlapped or flanked by our identified repeated sequences in the three genome assemblies analyzed. The size of the dot represents the number of genes contained in the gene set. A) GO analysis for genes found in GRCh37. B) GO analysis for genes found in GRCh38. C) GO analysis for genes found in T2T-CHM13.

Of the genes identified as potentially impacted by the identified repeats, 709, 721 and 691 genes in GRCh37, GRCh38 and T2T-CHM13, respectively, are annotated as disease associated in OMIM (Table 3). Among these are genes fulfilling dosage-sensitivity criteria that have been previously associated with genomic disorders, such as Bartter syndrome (*CLCNKB*, MIM #607364), Hajdu-Cheney syndrome (*NOTCH2*, MIM #102500), Ehlers-Danlos syndrome (*COL5A1*, MIM #130050), and Usher syndrome (*MYO7A*, MIM #276900) (Supplementary Table 2).

### Repeat overlap with genomic disorder regions and other reported structural variants and complex rearrangements

In order to evaluate the utility of our identified repeated sequence pairs for the study of more common genomic disorders, we cross-referenced the identified direct and inverted repeated sequences with regions known to be involved in the generation of recurrent and non-recurrent rearrangements. We observed clustering of repeats and the presence of larger repeat pairs flanking the reported deletion/duplication CNVs, consistent with known LCRs mediating recombination events in these regions (Supplementary Table 3, Supplementary Figures 2-4).

Additionally, we looked more broadly at the potential contribution of repeat pairs identified in this study to the generation of reported copy-number variants (CNVs) and structural variants observed in gnomAD (gnomAD SVs v2.1) (49). We observed a higher number of deletions flanked by direct repeats compared to duplications in all three assemblies, while for inversions we detected a small number of them flanked by inverted repeats across assemblies (Table 4). This observation could correlate with the ease of identifying each of these different types of variants by current methods. We also investigated CNVs flanked by repeats on only one side, as examples have been reported of some non-recurrent rearrangement breakpoints clustering within individual repeated sequences. Finally, we looked at the number of deletions, duplications, and inversions that overlap with the repeated sequences identified where we also observed an enrichment of deletions versus duplications or inversions across the three assemblies (Table 4). Given the scarcity of inversion variants available in public datasets with reported breakpoint junction information, we also looked at the overlap of inverted repeated sequences with previously reported inversions reported by Porubsky et al. mediated by mobile element insertions (MEIs) including LINE1 and *Alu* elements and SDs (7). Of the 65 reported MEI-mediated inversions, we found that 49 (75.38%) were overlapped by the identified repeated sequences, whilst 194 of 207 (93.71%) of the reported SD-mediated inversions overlapped with the repeats reported here (Supplementary Figure 5), supporting the usefulness of the dataset.

**Table 4.** Structural variants reported in gnomAD SVs V2.1 overlapped or flanked by identified repeated sequences.

| CNV | Flanking both sides |  |  | Flanking one side |  |  | Overlapping the CNV |  |  |
| --- | --- | --- | --- | --- | --- | --- | --- | --- | --- |
|  | GCRh37 | GCRh38 | T2T-CHM13 | GCRh37 | GCRh38 | T2T-CHM13 | GCRh37 | GCRh38 | T2T-CHM13 |
| Deletions | 4294<br>(69.4%) | 6779<br>(70.4%) | 4082<br>(70.3%) | 56023<br>(75.8%) | 55515<br>(76.3%) | 34460<br>(81.4%) | 10296<br>(74.2%) | 9817<br>(75.8%) | 7748<br>(81.09%) |
| Duplication | 1520<br>(24.5%) | 2430<br>(25.2%) | 1047<br>(19.58) | 16009<br>(21.6%) | 15745<br>(21.6%) | 6948<br>(16.4) | 3301<br>(23.8%) | 2880<br>(22.2%) | 1537<br>(16.09%) |
| Inversions | 373<br>(6.03%) | 410<br>(4.26%) | 219<br>(4.09%) | 1812<br>(2.45%) | 1461<br>(2.01%) | 907<br>(2.14) | 268<br>(1.93%) | 246<br>(1.9%) | 270<br>(2.83%) |

Finally, we also analyzed data of complex genomic rearrangements characterized by intermixed duplications and triplications of genomic segments in the *MECP2* region, previously reported in the literature (28,30). Notably, we identified the presence of inverted repeats at the breakpoint junctions of inverted triplications flanked by duplications, i.e. DUP-TRP/INV-DUP (Supplementary Figure 6) confirming previous observations and providing a catalog of genomic instability in at-risk regions.

## Discussion

Since the completion of the first draft of the human reference genome assembly (50,51), bioinformatic analyses of the human genomic sequence have led to the continued identification and refinement of genes, regulatory elements, enhancers, sequence motifs, and repetitive elements. However, the reference assembly focused primarily on the completion of the euchromatic regions of the human genome which are generally depleted of repeated sequences. Despite great improvements over the last two decades, the reference assembly still contains gaps and some regions have remained a challenge to map accurately, primarily due to the presence and enrichment of repetitive elements including tandem repeats, interspersed repeats, and segmental duplications (52,53). Recent assemblies utilizing long-read sequencing technologies offer the opportunity to explore the repetitive nature of the human genome across individuals and ancestries.

Genomic architectural features such as highly similar repeated sequences are important drivers of genome remodeling and evolution in the long term, but can also contribute in the short term to disease burden. Genomic disorders have been studied over more than two decades to understand the role of repetitive DNA sequences in remodeling the genome through recombination events that can result in clinically recognizable human disorders. These events can be facilitated by the presence of repetitive elements, including LCRs, SDs, SINEs and LINEs (54). Common chromosome deletion/duplication syndromes often involve rearranged genomic segments flanked by large LCR or SD structures that serve as recombination substrates. Overall, the instability and mutability of the genome are influenced by the presence of these repetitive sequences. Analyses of these genomic features, leveraging information observed *in vivo* and experimentally, can help predict regions of genomic instability that may result in genomic rearrangements. Studies that have aimed at performing this type of analyses focusing on potential substrates for NAHR events (12,42) have proven successful at demonstrating that predicted unstable regions indeed rearrange in human individuals (11,33). Conversely, retrospective analyses of complex genomic variants in human patients, such as Xq22 deletions that affect the *PLP1* gene resulting in a range of neurological disease traits in females, have shown the involvement of shorter inverted repeats in microhomology-mediated rearrangements (33,55).

In this study, we aimed to provide a genome-wide catalog of direct and inverted repeat sequences in the human genome that can potentially act as substrates for genomic rearrangements through different known mechanisms. This study investigates the distribution and genomic features of repeat pairs with high identity (80-100%) and with a minimum length of 200 bp. We analyzed their characteristics and distribution in three available human genome assemblies to evaluate the effect and impact of variable assemblies in the landscape of these repeats. Our findings show that a large portion of the human genome is potentially susceptible to genomic instability mediated by direct and inverted repeated sequences. No major differences were observed among the different assemblies, except for the newly resolved regions in the T2T-CHM13 assembly not present in the reference sequence. These regions had been difficult to resolve using the conventional sequencing methodologies utilized by the Human Genome Project due to their highly similar and repeat-rich architecture. They are mainly composed of SDs, ribosomal rRNA gene arrays, and satellite arrays that harbor unidentified sets of direct or inverted repeats previously overseen. Therefore, it is unsurprising, yet reassuring, that our analyses were able to identify a high density of repeat pairs and potentially unstable sequences within these regions.

The majority of the repeated pairs that we identified through the analyses reported here overlap repetitive elements in the human genome. Although previous analyses have looked at the distribution of highly identical repeats in the genome that may be substrates for NAHR (21,34,42), given the parameters that we used, we now recover repeat tracts formed by smaller repeated elements such as *Alus* that have been shown to be involved in microhomology-mediated genomic rearrangements through *Alu*-*Alu* mediated rearrangements (8,43). Further, we identified a significant fraction of protein-coding genes that are overlapped or flanked by the here reported repeated sequences. These are interesting because they represent “at risk” genes for potentially rearranging and leading to genomic disorders. Flanked genes are especially susceptible because if they are dosage-sensitive, they could be involved in genomic disorders through deletion or duplication events. Furthermore, in the case of overlapped genes, these could be disrupted by a genomic rearrangement potentially causing disease, especially if the genes are haploinsufficient. Interestingly, the majority of the identified “at risk” protein-coding genes are potentially dosage-sensitive, according to a recent report by Dong et al (45) that utilized RNA sequence data across different tissues to predict dosage sensitivity for genes in the human genome. Indeed, some known disease-associated dosage-sensitive genes such as *MYO7A* (Usher syndrome, MIM #276900), *CLCNKB* (Bartter syndrome, MIM #607364) and *SAMD9* (MIRAGE syndrome, MIM #617053) are recovered as flanked by the identified repeat pairs. The inversions reported by Porubsky et al. represent hotspots for morbid CNV formation associated with potential genomic disorder regions. These inversions change the relative orientation of the SD or MEI flanking elements. The new architecture of repeated elements surrounding the region might represent a protective or pre-mutational state for morbid CNV formation. Notably, these polymorphic inversions are often found to co-occur in regions linked to well-known genomic disorders, such as Williams-Beuren syndrome (WBS) and Smith-Magenis/Potocki-Lupski syndrome (SMS/PTLS). Understanding the distribution and characteristics of repeated sequences in the vicinity of these genes is crucial to gain insights into the potential causes of genomic instability and associated disorders. Our analyses also showed enrichment of repeats overlapping or flanking genes associated with sensory perception and immune response. CNVs have been previously reported to contribute to genetic variation in the human olfactory receptor (OR) repertoire (46,48). Our findings support the observations that the high abundance of repetitive elements in OR gene clusters contribute to genomic instability and variation at these loci. CNVs are known to impact genome evolution and adaptability by facilitating the expansion or contraction of gene families. Our findings contribute to the understanding of the potential impact of CNVs and repeated sequences on genome evolution in the context of olfactory receptors and other impacted biological processes.

Although the overall distribution of identified repeated sequences across human genome assemblies was very similar, it will be interesting to see if this holds similarly for other genomes as we start obtaining complete human genomes from individuals of diverse ancestries. The availability of long-read sequenced human genomes assembled *de novo* in a reference-free manner in the years to come offers the possibility to expand the landscape of human repeat variation and architecture. Analyses like ours will serve as good references to compare the genome-wide landscape and characteristics of these repeats across many human genomes in the near future. Overall, the results of this study provide a genome-wide map of potential sequences and sites that may serve as substrates for different recombination or replicative-associated mechanisms. These new datasets of direct and inverted repeated sequences in the three currently used human assemblies could help identify elements mediating novel copy-number variants and structural rearrangements that may have functional implications. These data may help uncover new disease-gene associations, facilitate molecular diagnosis, and offer further insights into genomic unstable regions and molecular mechanisms contributing to genome rearrangements.

## Methods

### Identification and consolidation of identical direct and inverted repeat pairs in the human genome

We performed bioinformatic analyses to identify repeated sequence pairs in direct and inverted orientations, with a minimum pairwise identity of 80% and a minimum length of 200bp, through self-alignment of each of the chromosomes of the three available human genome assemblies: GRCh37 (hg19), GRCh38 (hg38) and T2T-CHM13v2 using the LastZ 1.04.22 algorithm (56). The parameters used for minimum sequence identity and length were determined based on experimental data and observations reported in the literature by us and others for repeated sequences that can mediate genomic rearrangements through recombination mechanisms such as NAHR, or replication-based processes including MMBIR or FoSTeS.

For each assembly analyzed, after we obtained all the pairs of repeated sequences across the positive and negative strands of each chromosome representing the repeats on the same strand (direct repeats) or repeats on opposite strands (inverted repeats) we piped these results to a customized R code for downstream analyses. Due to the nature of LastZ, when an alignment exceeds a pre-specified minimum alignment score, the direct or inverted pair is reported even when the extension of the alignment is still possible before the homology drops below 80%. Therefore, we developed an algorithm to consolidate overlapping or nearby repeats in complex regions while still maintaining the percent identity between sequences at 80% or above (Supplementary Methods). The resulting datasets after repeat consolidation were used for downstream analyses, annotations, and comparisons.

### Repeat annotation and assembly comparisons

The resulting consolidated datasets for direct and inverted repeats of each of the three genome assemblies were cross-referenced with the coordinates of relevant genomic features such as segmental duplications, repetitive elements, protein-coding genes, and reported structural variants.

We used CrossMap (57) to perform coordinate liftover between the repeats obtained in each of the three genome assemblies analyzed for comparisons. This allowed us to check for sequence overlap between elements in the different assemblies and compare the genome-wide distributions of repeats between and across human genome assemblies.

We looked for overlap between repeated sequences identified here and known repetitive elements, such as SINEs, LINEs, SDs, and satellite DNA. Therefore, we compared the direct and inverted repeat datasets for all three assemblies with the RepeatMaskerViz dataset to identify the type of repetitive elements that overlapped our repeated sequences in each assembly. Additionally, we cross-referenced our repeats datasets with RefSeq protein-coding genes to identify genes overlapped or flanked by repeated sequences, OMIM annotations for genes associated with human diseases and the Dong et al (45) predicted dosage-sensitive gene list. For overlap, we looked at any repeats overlapping genes by at least 30%, whereas for genes flanked by pairs of repeated sequence we focused on pairs with features compatible with potential for NAHR, mainly >90% sequence identity and up to a distance of 100 kb upstream or downstream.

### Ontology analysis

The topGO package in R was utilized for functional enrichment analysis, with the org.Hs.eg.db annotation package providing Gene Ontology (GO) terms. The gene list used for the ontology analysis was obtained from the previous downstream analysis described before to identify genes overlapped and/or flanked by the identified repeated sequences. GO enrichment analysis of the input gene lists was performed using the runTest function in Gene Ontology, employing Fisher’s exact test to determine the significance of gene set overrepresentation in specific GO terms. The “topGO” package’s weight algorithm assigned weights to GO terms based on their specificity, facilitating the determination of the number of genes annotated to each term.

### Overlap with experimentally validated reported rearrangements

To evaluate the utility of our bioinformatically identified repeated sequences, we looked at the overlap or flanking of experimentally validated structural variants and genomic rearrangements with our datasets. We bioinformatically cross-referenced the coordinates of our repeats in the different assemblies with known genomic disorders (25), inversions reported by Porubsky et al (7), and the gnomAD SVs v2.1 dataset (49). The corresponding available datasets were obtained and we used bedtools to intersect the coordinates of our direct and inverted repeats datasets with the corresponding regions. Most reported datasets were provided in GRCh37 coordinates, so we used CrossMap (57) to perform coordinate liftover to the GRCh38 and T2T-CHM13 assemblies. For flanking regions, coordinates were obtained using bedtools flank, with a maximum distance of 100 kb. The flanking regions were overlapped with the set of direct repeats provided in the study, and downstream analyses were performed to filter and keep pairs of direct repeats with a homology above 80% flanking on both sides. Additionally, repeats that were flanking only one side of the CNV and those overlapping the entire CNV were also identified (Table 4).

To visualize repeats and features in genomic regions of interest, we uploaded our generated direct and inverted repeat tracks to the UCSC browser and looked at other tracks of interest such as pathogenic deletion and duplications from ClinVar (Supplementary Figure 2-4), OMIM genes, Repeat Masker, etc.

## Acknowledgments

The authors wish to thank Luis Alberto Aguilar, the Laboratorio Nacional de Visualización Científica Avanzada (LAVIS) and the International Laboratory for Human Genome Research of UNAM, for computational resources and support to perform the analyses reported here. C.M.B.C. is supported by the National Institute of General Medical Sciences (NIGMS R01 GM132589).

